# Longitudinal linear-epitope profiling of SLE identifies novel antigens associated with disease and disease activity

**DOI:** 10.64898/2026.09.29.755420

**Authors:** Charlotte Hurabielle, Yutong V. Wang, Bryan Castillo-Rojas, Chung-Yu Wang, Colette Caspar, James Asaki, Monica Dayao, Max Calcoen, Eric D. Merrill, Aaron Bodansky, Arielle Klepper, Elizabeth N. Chung, Caroline Vissers, Kelsey C. Zorn, Mark S. Anderson, Michael R. Wilson, Peter Izmirly, Jill Buyon, Jinoos Yazdany, Maria Dall’Era, Chun J. Ye, Joseph L. DeRisi

**Affiliations:** Division of Rheumatology, Department of Medicine, University of California, San Francisco, San Francisco, CA, USA; Institute for Human Genetics, University of California, San Francisco, San Francisco, CA, USA; Biohub, San Francisco, CA 94158, USA; Department of Biochemistry & Biophysics, University of California, San Francisco, San Francisco, CA, USA; Weill Institute for Neurosciences, Department of Neurology, University of California, San Francisco, San Francisco, CA, USA; Gladstone-UCSF Institute of Genomic Immunology, San Francisco, CA, USA; Department of Dermatology, University of California, San Francisco, San Francisco, CA, USA; Department of Dermatology, Veterans Affairs Medical Center, San Francisco, CA, USA; Department of Pediatric Critical Medicine, University of California San Francisco; San Francisco, CA; Child Health Advanced Molecular Phenotyping (CHAMP) Program, University of California San Francisco; San Francisco, CA; Department of Medicine, University of California, San Francisco, San Francisco, CA, USA; UCSF Liver Center, University of California, San Francisco, San Francisco, CA, USA; Tetrad Graduate Program, University of California, San Francisco, San Francisco, CA, USA; Medical Scientist Training Program, University of California, San Francisco, San Francisco, CA, USA; Immunectra, Inc; Diabetes Center, UCSF; Division of Rheumatology, Department of Medicine, NYU Grossman School of Medicine, New York, NY, USA; Division of Rheumatology, Zuckerberg San Francisco General Hospital, San Francisco, CA, USA; Department of Research and Early Development: Translational Genomics and Proteomics Center, Genentech, San Francisco, California, USA; Parker Institute for Cancer Immunotherapy, San Francisco, CA, USA; Arc Institute, Palo Alto, CA, USA; Bakar Aging Research Institute (BARI), University of California, San Francisco, San Francisco, CA, USA; Department of Bioengineering and Therapeutic Sciences, University of California, San Francisco, San Francisco, CA, USA; Department of Epidemiology and Biostatistics, University of California, San Francisco, San Francisco, CA, USA

## Abstract

Systemic lupus erythematosus (SLE) is characterized by diverse autoantibodies, yet established biomarkers capture only a fraction of autoreactivity. We profiled 1,268 plasma samples from 695 patients with SLE and 301 healthy controls using phage immunoprecipitation sequencing (PhIP-seq). Machine learning identified a reproducible autoreactivity signature that discriminated SLE from controls across two validation cohorts. SLE autoreactivities were enriched for RNA-binding proteins, while targets containing Basic Acidic Dipeptide repeat domains were associated with high disease activity. A YTHDF2-derived peptide was the most prevalent shared autoreactivity, occurring in ∼25% of patients and exhibiting sequence similarity to Epstein-Barr virus (EBV) lytic switch protein BZLF1. Competition assays demonstrated higher-affinity binding to the viral peptide, consistent with EBV-associated molecular mimicry. Orthogonal validation using a 10-peptide Luminex panel strongly discriminated SLE from healthy controls. These findings expand the SLE autoantibody landscape, identify disease activity associated autoreactivities and insight linking EBV-driven immunity to epitope-specific autoimmunity in lupus.

## Introduction

Systemic lupus erythematosus (SLE) is the prototypical systemic autoimmune disease defined by the production of autoantibodies directed against a broad spectrum of self-antigens^1^. These humoral immune responses contribute directly to disease pathogenesis through immune complex deposition, complement activation, and tissue inflammation across multiple organ systems. Despite marked clinical heterogeneity, the presence of antinuclear antibodies (ANAs) is a defining feature of SLE and remains central to disease classification and diagnosis^2^. However, conventional serologic assays, including ANA, anti-dsDNA, and extractable nuclear antigen (ENA) testing, capture only a limited subset of autoreactivities and provide incomplete insight into the broader organization and origins of the autoreactive immune response^3^.

Recent therapeutic advances have further underscored the central role of B cells in SLE pathogenesis. Targeted depletion of B and plasma cells, including anti-CD20 monoclonal antibodies and BAFF inhibition, has demonstrated clinical benefit in subsets of patients with lupus^4–8^. More recently, CD19 and BCMA CAR T cell therapies have induced durable drug-free remissions in patients with severe, refractory disease^9–13^. The promising benefit attained by B cell depletion further underscores the importance of both B cells and the autoantibodies they produce.

Most current knowledge of lupus autoreactivity derives from targeted assays focused on a limited number of canonical antigens. Previous discovery efforts, including immunoprecipitation, immunoblotting, expression library screening, and, more recently, protein microarrays^14–16^ were justifiably designed around full length, folded proteins or native macromolecular complexes. As a result, these techniques were not optimized for the systematic discovery of linear peptide epitopes, despite growing evidence that the human autoreactome contains numerous such determinants. Given the complexity and heterogeneity of SLE, we hypothesized that unbiased mapping of autoreactive linear epitopes across patients would yield a distinct set of new and complementary features that both expand our understanding of the humoral targets of SLE and clinically stratify patients^17^. Such an approach may also provide insight into the longstanding question regarding the origins of lupus autoreactivity. Among environmental factors implicated in SLE, Epstein Barr virus (EBV) has emerged as one of the strongest and most reproducible associations^18^. Recent studies have provided mechanistic evidence that EBV infection can directly reprogram autoreactive B cells in lupus, strengthening longstanding models linking antiviral immunity to systemic autoimmunity^19^. In parallel, multiple studies have demonstrated a near-universal association between EBV infection and multiple sclerosis (MS), and have further reinforced interest in molecular mimicry as a mechanism by which tolerance is broken against a highly prevalent latent infection resulting in the development of autoimmunity^20–26^. Unlike MS, the epitope-specific links between antiviral immunity and lupus autoreactivity remain less well defined.

Here, phage immunoprecipitation and sequencing (PhIP-seq)^27^ was used to profile autoantibody responses against the linear human peptidome across three separate SLE cohorts. We identify a reproducible autoreactive signature enriched for RNA-binding proteins and proteins containing basic–acidic dipeptide (BAD) domains, which have been associated with biomolecular condensate formation. We further define a shared autoreactive epitope within YTHDF2 that exhibits sequence homology and functional cross-reactivity with the key EBV lytic switch protein BZLF1, providing mechanistic support for a model in which antiviral humoral immunity may promote lupus-associated autoreactivity through molecular mimicry and epitope spreading. Despite the complexity and breadth of the SLE autoreactive signature, we further demonstrate that this can be compressed into a simple, 10-peptide on Luminex bead assay that shows robust SLE classification and detection capability in a validation cohort.

## RESULTS

### Proteome-wide interrogation of 890 SLE samples, compared to 208 healthy controls yields a distinct and robust humoral immune profile

The overall approach, including analysis and validation, for this study is illustrated in **Figure 1A**. To investigate and define the serological landscape of SLE at the linear peptide resolution, a discovery cohort (cohort #1), comprised of a total of 890 samples from 475 patients was obtained from the California Lupus Epidemiology Study (CLUES) (**Table 1**, Methods) in addition to 208 healthy control (HC) samples. Plasma from each of the samples was used to iteratively immunoprecipitate a previously validated human proteome phage display library designed with 731,724 tiled 49-mer peptides spanning the human proteome^17,23,28–31^.

**Figure 1.**
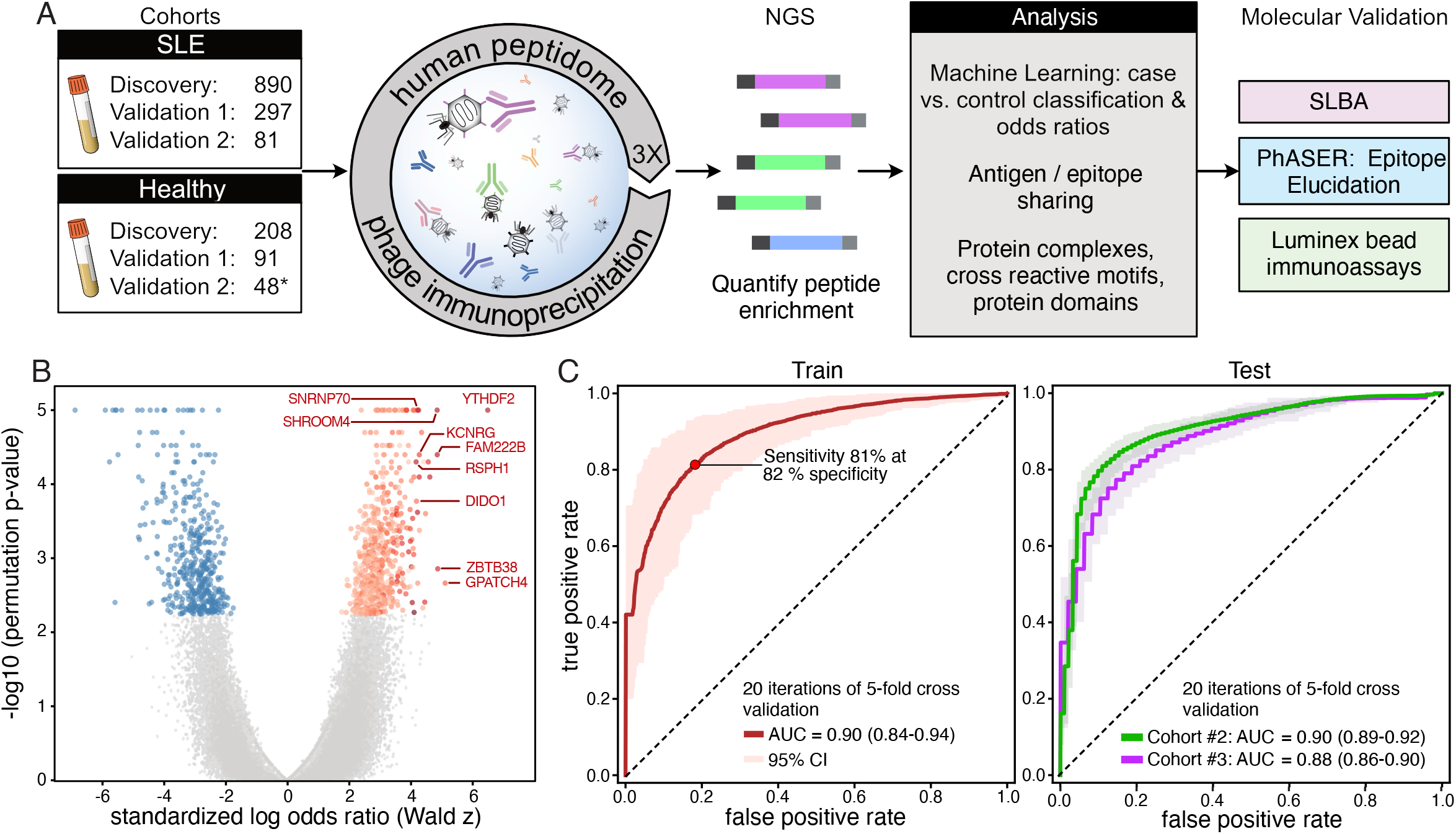
Large-scale PhIP-seq profiling identifies a robust and generalizable humoral immune signature in SLE. A. Overview of the PhIP-seq pipeline and methods used for analysis and validation comparing 3 separate cohorts of SLE samples (total n=1,268) and healthy controls (total n=301). 46 healthy controls in cohort 3 were also sampled in cohort 1 and are counted once in the total (see Methods). B. Protein level differential autoantibody reactivity between SLE donors and healthy controls, colored when Benjamini-Hochberg FDR < 0.1, in red for SLE-enriched, blue for HC-enriched. Red intensity indicates the fraction of SLE donors with a peptide Z-score at or above 4.5. C. Receiver Operating Characteristic curves for the logistic regression classifier of SLE versus healthy control, trained in the discovery cohort (train, left) and evaluated in two independent validation cohorts (test, right) with 95% confidence interval.

**Table 1.** Cohort demographic and clinical characteristics.

|  | Cohort 1 |  |  | Cohort 2 |  |  | Cohort 3 |  |  |
| --- | --- | --- | --- | --- | --- | --- | --- | --- | --- |
|  | SLE samples | SLE at inclusion | healthy controls | SLE samples | SLE at inclusion | healthy controls | SLE samples | SLE at inclusion | healthy controls |
| n | 890 | 475 | 208 | 297 | 161 | 91 | 81 | 59 | 48* |
| female | 787 (88.4%) | 424 (89.3%) | 150 (72.1%) | 270 (90.9%) | 144 (89.4%) | 47 (51.6%) | 75 (92.6%) | 54 (91.5%) | 43 (89.6%) |
| median age (years) | 39 [29.2; 53] | 41 [30; 54] | 30 [24-46] | 31 [25.0;41.2] | 30 [24; 38] | 46 [31.5; 54.0] | 34 [23.0;39.0] | 28 [23.0;39.0] |  |
| age at diagnosis (years) | 25 [19; 34.8] | 26 [20; 35] |  | 21.1 [17.9; 29.4] | 21 [17.9; 28.7] |  |  |  |  |
| median disease duration (years) | 13 [6; 21] | 12 [6; 21] |  | 8.8 [5.0; 15.1] | 8.5 [4.7; 14.5] |  |  |  |  |
| **race/ethnicity** |  |  |  |  |  |  |  |  |  |
| east asian | 301 (33.8%) | 167 (35.2%) | 38 (18.3%) | 27 (9.1%) | 10 (6.2%) | 3 (3.3%) | 10 (12.3%) | 6 (10.2%) |  |
| hispanic/latino | 242 (27.1%) | 117 (24.6%) | 14 (6.7%) | 37 (12.5%) | 17 (10.6%) | 13 (14.3%) | 42 (51.9%) | 30 (50.8%) |  |
| white | 208 (23.4%) | 120 (25.3%) | 25 (12.0%) | 94 (31.6%) | 64 (39.8%) | 69 (75.8%) | 11 (13.6%) | 10 (16.9%) |  |
| african american | 115 (12.9%) | 58 (12.2%) | 5 (2.4%) | 100 (33.7%) | 63 (39.1%) | 1 (1.1%) | 11 (13.6%) | 6 (10.2%) |  |
| other | 24 (2.7%) | 13 (2.7%) | 9 (4.3%) | 6 (2.0%) | 7 (4.3%) | 5 (5.5%) | 7 (8.6%) | 7 (11.9%) |  |
| unknown |  |  | 107 (51.4%) | 33 (11.1%) | 4 (2.5%) |  |  |  |  |
| median sledai score | 4 [2; 8] | 4 [0; 6] | N/A | 6 [2.0;12.0] | 11.0 (7.2–15.5) |  | 2 [0.0;8.0] | 2 [0.0;14.0] |  |
| none (0) | 196 (22%) | 119 (25%) |  | 10 | 0 |  | 29 (35.8%) | 23 (39.0%) |  |
| mild (1-5) | 384 (43.1%) | 206 (43.4%) |  | 39 | 6 |  | 30 (37.0%) | 16 (27.1%) |  |
| moderate (6-10) | 179 (20.1%) | 75 (15.8%) |  | 23 | 13 |  | 4 (4.9%) | 3 (5.1%) |  |
| high (≥11) | 130 (14.6%) | 75 (15.8%) |  | 27 | 19 |  | 18 (22.2%) | 17 (28.8%) |  |
| unknown | 1 (0.1%) |  |  | 198 | 123 |  |  |  |  |
| **active disease category (percentage among patients with data)** |  |  |  |  |  |  |  |  |  |
| elevated dsDNA Ab | 479 (53.8%) | 247 (52%) |  | 78 (78.8%) | 33 (86.8%) |  | 43 (53.1%) | 29 (49.2%) |  |
| low complement | 399 (44.8%) | 194 (40.8%) |  | 53 (53.5%) | 30 (78.9%) |  | 39 (48.1%) | 31 (52.5%) |  |
| nephritis† | 193 (21.7%) | 92 (19.4%) |  | 39 (39%) | 28 (73.7%) |  | 20 (24.7%) | 19 (32.2%) |  |
| rash | 157 (17.6%) | 71 (14.9%) |  | 15 (15%) | 9 (23.7%) |  | 7 (8.6%) | 7 (11.9%) |  |
| arthritis | 131 (14.7%) | 66 (13.9%) |  | 8 (8%) | 6 (15.8%) |  | 4 (4.9%) | 3 (5.1%) |  |
| leukopenia | 72 (8.1%) | 41 (8.6%) |  | 6 (6.1%) | 4 (10.5%) |  | 8 (9.9%) | 3 (5.1%) |  |
| alopecia | 59 (6.6%) | 28 (5.9%) |  | 19 (19%) | 10 (26.3%) |  | 4 (4.9%) | 4 (6.8%) |  |
| mucosal ulcers | 48 (5.4%) | 32 (6.7%) |  | 5 (5%) | 3 (7.9%) |  | 1 (1.2%) | 1 (1.7%) |  |
| thrombocytopenia | 43 (4.8%) | 24 (5.1%) |  | 0 (0%) | 0 (0%) |  | 0 (0%) | 0 (0%) |  |
| serositis | 33 (3.7%) | 20 (4.2%) |  | 2 (2%) | 2 (5.3%) |  | 3 (3.7%) | 3 (5.1%) |  |
| neuro/psychiatric involvement | 16 (1.8%) | 11 (2.3%) |  | 0 (0%) | 0 (0%) |  | 0 (0%) | 0 (0%) |  |
| fever | 16 (1.8%) | 10 (2.1%) |  | 0 (0%) | 0 (0%) |  | 1 (1.2%) | 1 (1.7%) |  |
| vasculitis | 12 (1.3%) | 7 (1.5%) |  | 5 (5%) | 2 (5.3%) |  | 0 (0%) | 0 (0%) |  |
| myositis | 2 (0.2%) | 1 (0.2%) |  | 2 (2%) | 2 (5.3%) |  | 0 (0%) | 0 (0%) |  |
| **lupus medication use, n (%)** | **761 (85.5%)** | **385 (81%)** |  |  |  |  | **68 (84%)** | **52 (88.1%)** |  |
| Hydroxychloroquine | 644 (72.4%) | 326 (68.6%) |  |  |  |  | 60 (74%) | 47 (79.7%) |  |
| Steroids | 495 (55.6%) | 242 (50.9%) |  | 239 (80.5%) | 139 (86.3%) |  | 28 (34.5%) | 22 (37.3%) |  |
| Mycophenolate | 290 (33%) | 139 (29.3%) |  |  |  |  | 19 (23.5%) | 12 (20.3%) |  |
| Azathioprine | 79 (8.9%) | 39 (8.2%) |  |  |  |  | 0 (0%) | 0 (0%) |  |
| Methotrexate | 58 (6.5%) | 26 (5.5%) |  |  |  |  | 6 (7.4%) | 5 (8.5%) |  |
| Cyclophosphamide IV | 33 (3.7%) | 14 (2.9%) |  |  |  |  | 3 (3.7%) | 2 (3.4%) |  |
| Rituximab or Obinutuzumab | 25 (2.8%) | 6 (1.3%) |  |  |  |  | 6 (7.4%) | 4 (6.8%) |  |
| CNI | 38 (4.3%) | 18 (3.8%) |  |  |  |  | 3 (3.7%) | 2 (3.4%) |  |
| Tacrolimus | 24 (2.7%) | 15 (3.2%) |  |  |  |  | 3 (3.7%) | 2 (3.4%) |  |
| Cyclosporine | 13 (1.5%) | 3 (0.6%) |  |  |  |  |  |  |  |
| Voclosporine | 1 (0.1%) | 0 (0%) |  |  |  |  |  |  |  |
| Belimumab | 14 (1.6%) | 7 (1.5%) |  |  |  |  | 2 (2.5%) | 1 (1.7%) |  |
| Abatacept | 12 (1.3%) | 8 (1.7%) |  |  |  |  |  |  |  |
| Leflunomide | 11 (1.2%) | 8 (1.7%) |  |  |  |  |  |  |  |
| IVIG | 4 (0.4%) | 4 (0.8%) |  |  |  |  |  |  |  |
| Anifrolumab | 1 (0.1%) | 0 (0%) |  |  |  |  | 2 (2.5%) | 2 (3.4%) |  |

† Any positive among sledai\_hematuria, sledai\_proteinuria, sledai\_pyuria, sledai\_urinarycasts
\* 46 of the 48 healthy controls originated from the cohort #1, see methods
n; number; SLE, systemic lupus erythematosus; dsDNA Ab, anti-double-stranded DNA antibodies; CNI, calcineurin inhibitors; IVIG: intravenous immunoglobulins.

To assess how the autoantibody profiles for SLE cases differed from controls globally, per-peptide enrichment statistics and false discovery rates were computed for all peptides in this human peptidome library. Consistent with prior PhIP-seq studies, we found that each individual in our cohort harbored a unique constellation of personal peptide autoreactivities^32,33^; however, the overall magnitude of autoreactivity was significantly higher in SLE: across all 360,920 peptides with more than 10 reads across the whole study population, the per-sample mean reactivity z-score was shifted higher in SLE than HC (Mann-Whitney p = 2.6×10^-14^, **Suppl. Fig. 1A**).

To assess differences in protein-level autoreactive enrichment between SLE and healthy controls, a permutation test using the combined-peptide results was applied and proteins were ranked and plotted by the signed standardized log odds ratio (Wald z), with the sign giving the direction of enrichment (Methods, **Fig 1B**.) This approach identified 647 autoreactive proteins enriched in SLE plasma samples compared to healthy control plasma samples, suggesting extensive sharing of autoreactive target antigens between patients. We hypothesized that profiling a linear peptide representation of the proteome would surface a set of previously unrecognized, largely non-overlapping antigens with those previously discovered by methods based on whole proteins. Consistent with this notion, the antigens identified here are almost entirely distinct from previously discovered autoantigens used in clinical practice. Exceptions include the classical SLE associated autoantigen SNRNP70, which is a core component of the U1 small nuclear ribonucleoprotein, associates directly with the Sm proteins, and is detected in up to 20-30% of SLE patients^34^. Here, SNRNP70 presents with a standardized log odds ratio (log(OR)) of 4.21 and a permutation p-value of 1.00×10^-5^, indicative of one of the most significant hits in this data set. However, autoreactive targets such as YTHDF2 (standardized log(OR) = 6.48, p-val < 10^-5^), were even more prevalent, yet are previously unreported in SLE. Consistent with our thesis, these results significantly expand the spectrum of SLE-associated autoreactivities beyond previously discovered clinical serologies and define a broader antigenic landscape than was previously appreciated.

### An autoantibody classifier discriminates SLE from controls and generalizes to additional cohorts of patients

To further define and evaluate the SLE-associated autoreactive signature, a regularized logistic regression classifier (Elastic Net) was trained using the discovery cohort. To prevent information leakage from repeated sampling of the same individual, all samples from a given patient were assigned to the same cross-validation fold, and class imbalance was addressed by class weighting. Across 20 repeated five-fold cross-validation runs, the classifier achieved a mean area under the receiver operating characteristic curve (AUROC) of 0.90 (95% CI, 0.84-0.94), with 81% sensitivity at 82% specificity (**Fig. 1C, left**).

To assess generalizability, we applied the trained classifier to two independent external cohorts comprising 297 SLE and 91 healthy controls (cohort #2) and 81 SLE and 48 healthy controls (cohort #3). Importantly, none of these samples were used during model training or feature selection. The classifier demonstrated comparable performance in both validation cohorts, achieving AUCs of 0.90 (95% CI, 0.89-0.92) and 0.88 (95% CI, 0.86-0.90) respectively (**Fig. 1C, right**), indicating robust generalizability. Model performance was largely preserved when restricting features to the highest-ranked peptides. Indeed, using only the top 10 peptides preserved strong discrimination (AUC 0.77 and 0.81 in validation cohorts 2 and 3, respectively). Performance remained high even when reduced to the top 5 peptides YTHDF2, RSPH1, NT5C1A, SNRNP70, DDX17; AUC 0.72 and 0.79) or even using a single peptide (YTHDF2, AUC 0.66 and 0.75; **Suppl. Fig. 1B-C**), consistent with the notion that a small number of shared linear peptide autoreactivities captures meaningful predictive power.

### Co-occurring recognition across multiple targets within individuals suggest a shared antigenic epitope

To assess the extent of autoreactive antigen sharing amongst patients, the 20 most significantly enriched SLE associated antigens were extracted from cohort #1 (FDR < 0.1, ranked by signed standardized log(OR) (Wald z)), and compared to the other two cohorts and healthy controls (**Fig 2A**). Here, we found a complex pattern of sharing, with few patients sharing exactly the same autoreactivity over these 20 targets. Nevertheless, autoreactivity across this antigen set was substantially more prevalent among patients with SLE than healthy controls, which showed only sporadic enrichment that rarely exceeded a generous z-score threshold of 2.5. In several instances, such as YTHDF2, RSPH1, and SNRNP70, overlapping peptides were consistently enriched, suggesting autoantibodies for epitopes specifically within the overlap region of the peptide tiles. As noted previously, with the exception of SNRNP70, these top 20 proteins represent new features of the humoral autoreactivities in SLE; however, we note numerous instances of adjacency to complexes or molecular processes associated with previously described SLE autoantigens. These include histone 3 variants H3F3A/C (which encode nearly identical proteins), other chromatin associated proteins such as L3MBTL3, ZBTB38, DIDO1, and RNA processing proteins such as SNRNP70, GPATCH4, and YTHDF2.

**Figure 2.**
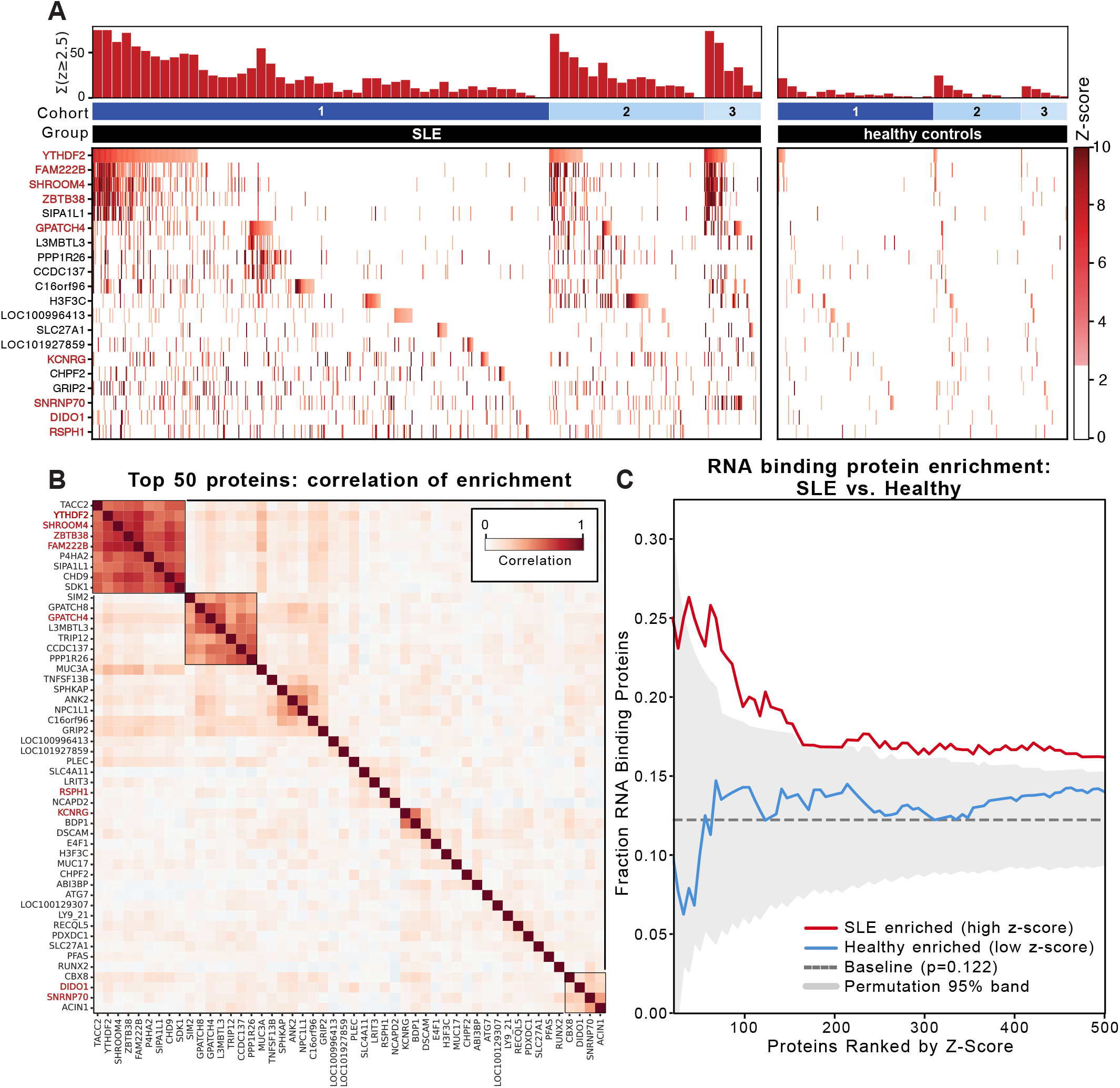
Enrichment of RNA binding proteins amongst SLE-specific autoantibodies. A. Heatmap of significantly enriched peptides in SLE (left) compared with healthy controls (right) across the training cohort (1) and validation cohorts (2 and 3). Rows are the top 20 SLE-enriched proteins from Fig 1B (Benjamini-Hochberg FDR < 0.1, ranked by standardized log odds ratio). Each column represents an individual sample (each individual only represented once), and each cell is colored according to its peptide enrichment Z-score relative to healthy controls; Z-scores < 2.5 are shown in white. The histogram above the heatmap depicts the total number of peptide enrichments with Z-scores ≥ 2.5 summed across all displayed peptides within consecutive bins of 10 samples, reset for each cohort change. B. Pearson correlation of co-occurrence among the top 50 peptides enriched in SLE. C. Fraction of RNA binding proteins (RBPs) among top K proteins, for K from 1 to 500, enriched in SLE (red) and healthy controls (blue) with grey band showing 95% confidence interval with permutation.

Expanding to the 50 highest ranked SLE enriched autoreactive proteins, a pairwise correlation matrix of per-donor z-scores was constructed to specifically detect co-occurrence patterns in the data (**Fig 2B**). Two major co-occurrence blocks were immediately evident, followed by several weaker correlations among patients. The first contained 9 proteins, including TACC2, YTHDF2, SHROOM4, and ZBTB38, that were highly correlated (mean Pearson r=0.65, permutation p-value < 10^-5^), suggesting the presence of shared autoreactive sequence features. Sequence analysis of peptides within the dominant co-occurring cluster revealed a conserved PxNAY motif present across multiple enriched targets. This motif was embedded within the overlapping YTHDF2 peptides and was also detected in three of the other highly co-enriched antigens: SDK1, CHD9, and P4HA2. Sequence inspection of the second cluster of co-occurring autoreactivities (r=0.50, p-value < 10^-4^), comprising peptides derived from SIM2, GPATCH4, GPATCH8, L3MBTL3, TRIP12, CCDC137, and PPP1R26, revealed a strong enrichment for basic residues, particularly recurrent di-arginine (RR), lysine/arginine (K/R) pairs, and arginine/serine (RS/SR) dipeptides. The enriched peptides (49aa) in this second cluster have a mean isoelectric point (pI) of 9.4, compared to 7.1 for randomly chosen peptides of the same length (p= 5.2×10^-3^), and a lysine/arginine content of 21% vs. 11% for randomly chosen peptides (p=1.6×10^-4^). These low complexity sequences are typical of nucleic acid binding proteins^35^ and intrinsically disordered regions. The enriched autoreactivity may be the consequence of shared physicochemical features, rather than a specific binding motif.

### Enrichment of RNA-binding proteins amongst SLE-specific autoantibodies

Protein-level enrichment analysis revealed a significant overrepresentation of RNA-binding proteins (RBPs) among SLE-enriched targets (**Fig. 2C**). Proteins annotated with RNA-binding activity, including SNRNP70, YTHDF2, FIP1L1, DIDO1, GPATCH4, YTHDC1, were disproportionately represented among top-ranked proteins (top 500) compared to expectation under permutation (p < 2×10^-3^, see Methods). This enrichment was most pronounced among the highest-effect-size proteins and diminished with inclusion of lower-ranked targets, consistent with concentrated pathway-level bias. While previous work has identified XIST-associated proteins as targets of autoantibodies in SLE and other female-biased autoimmune diseases^36^, and YTHDF2 has been reported to immunoprecipitate XIST RNA with increased levels of XIST with YTHDF2 knowckdown^37^, the collection of RNA binding proteins described here was otherwise not significantly biased toward XIST RNA binding proteins as a class (p=0.32). Together, these findings extend the canonical concept of ENAs by demonstrating preferential humoral targeting of RBPs in SLE.

### High disease activity is associated with autoantibodies against basic-acidic dipeptide repeats

Although the autoantibody landscape clearly distinguishes SLE from healthy controls, it was unclear whether the breadth and specificity of humoral autoreactivity varied with clinical disease activity. Anti-dsDNA IgG is typically used clinically to track disease activity longitudinally, however it is an imperfect indicator of disease state, since titers often stay elevated in patients without clinical manifestations^38^. To determine whether autoantibody profiles in this assay distinguish active from quiescent disease, we analyzed patients at extremes of clinical activity. A penalized logistic regression classifier distinguished patients with high disease activity (SLEDAI ≥11) from those with inactive disease (SLEDAI = 0) under donor-grouped cross-validation (**Fig. 3A**), and this performance was reproduced in the two validation cohorts.

**Figure 3.**
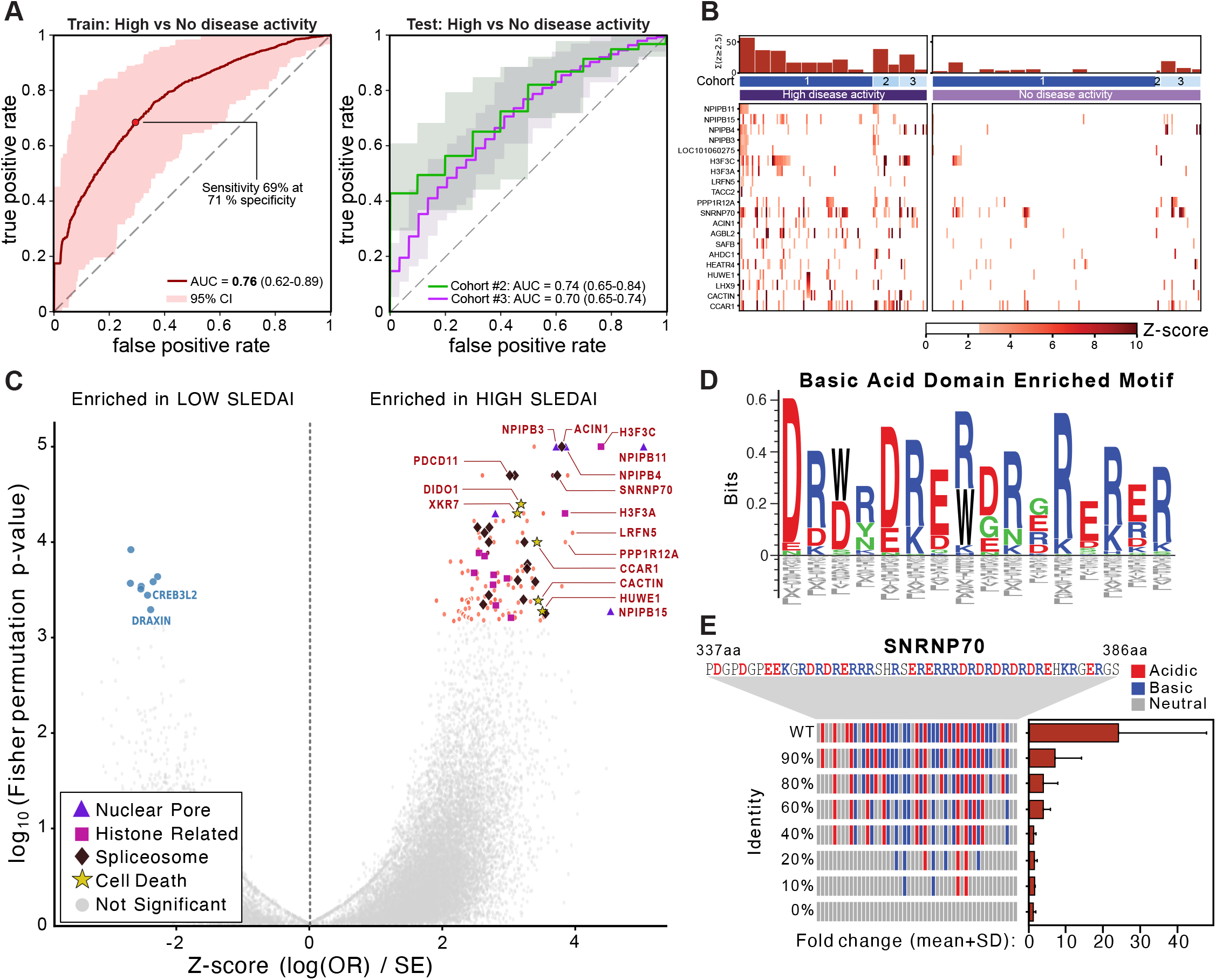
Autoantibodies against basic acidic dipeptide (BAD) domains distinguishes SLE disease activity. A. Receiver Operating Characteristic (ROC) curves for the logistic regression classifier of high disease activity (SLEDAI ≥ 11) as compared to no disease activity (SLEDAI=0) among SLE donors, trained in the discovery cohort (train, left) and evaluated in two independent validation cohorts (test, right) with 95% confidence interval. B. Heatmap of significantly enriched peptides in high disease activity (SLEDAI ≥ 11) (left) as compared to no disease activity (SLEDAI=0) (right), in the training (1) and validation cohorts (2 and 3). Rows are the most significantly enriched proteins in high versus no disease activity from Fig 3C (Benjamini-Hochberg FDR < 0.1, ranked by standardized log odds ratio). Each column represents an individual sample, and each cell is colored according to its peptide enrichment Z-score relative to healthy controls; Z-scores < 2.5 are shown in white. The histogram above the heatmap depicts the total number of peptide enrichments with Z-scores ≥ 2.5 summed across all displayed peptides within consecutive bins of 10 samples, reset for each cohort change. C. Volcano plot comparing high disease activity with no disease activity autoreactivities, highlighting different groups of proteins: nuclear pore, histone related, spliceosome, and cell death related. D. Sequence logo revealing enriched basic acidic dipeptide (BAD) domain across enriched peptides in patients with high disease activity. E. Ligand binding assay of variants of peptide derived from SNRNP70 starting with original sequence (wild type, WT), followed by sequential decrease in the proportion of basic and acidic peptides.

Protein-level enrichment analysis revealed a subset of autoantibody targets preferentially associated with high disease activity (**Fig. 3B,C**). Among the most strongly enriched targets were multiple RBPs and nuclear factors, suggesting that high disease activity disease is accompanied by intensified targeting of chromatin, spliceosome, and nuclear pore complex or envelope. These include factors like SNRNP70, and histone H3F3A/C, but also multiple members of the nuclear pore complex-interacting family members, such as NPIPB3, NPIPB4, NPIPB11, and NPIPB15, which feature prominent intrinsically disordered regions. Together, this pattern may reflect an increased load of cellular debris and associated protein condensates in the wake of cytotoxicity.

Using the sequences of all the peptides (n=123) significantly enriched in high disease activity, we looked for enriched motifs using MEME^39^, which revealed a subset sharing a sequence corresponding to the basic acidic dipeptide (BAD) repeat domain (E value = 5.9×10^-17^) (**Fig 3D**).

BAD domains are intrinsically disordered, low-complexity regions defined by highly repetitive alternating basic (Arg/Lys) and acidic (Asp/Glu) residues. They are a known structural motif present in the C-terminal low-complexity region of SNRNP70 (U1-70K) and were shared across 9 peptides significantly enriched in high disease activity patient samples (SNRNP70, NELFE, RBM25, ACIN1, DIDO1, ZC3H13, PRPF38B, CCAR1, SRRT). To probe autoreactivity to this motif, we performed systematic alanine disruption of the BAD motif: sequential substitution of the basic-acidic residues from SNRNP70 with alanine (10– 100% of motif positions mutated) led to progressive loss of antibody reactivity in patient plasma (n=5) using a split-luciferase binding assay (SLBA) (**Fig. 3E**, see Methods). More than 50% of the residues in this 49aa sequence were required to be mutated before all autoreactivity was abolished. This mutation-dependent reduction in signal supports the hypothesis that the basic-acidic charge alternation itself is determinant for autoantibody binding.

Charged multivalent interaction motifs in RBPs have been implicated in biomolecular condensate assembly and phase separation^40^. Specifically for SNRNP70, the BAD domain has been shown to drive liquid-liquid phase separation and higher-order self-assembly in vitro and in nuclear condensates^41^. We thus asked whether the proteins preferentially targeted in active SLE exhibit an increased propensity for condensate localization. To address this, we computed PICNIC scores, a machine-learning derived metric that outputs the predicted probability of a protein being a condensate member, where higher scores indicate a higher predicted probability^42^. Proteins differentially enriched in the high-disease-activity group displayed significantly higher PICNIC scores (median 0.63 vs 0.44, p-val <1×10^-5^, see Methods) than randomly selected proteins (**Suppl. Fig. 3 A)**. These results are consistent with the notion that the SLE disease process drives autoreactivity toward proteins with condensate propensity.

### Identification of YTHDF2 epitope reveals homology with EBV BZLF1

Peptide sequences derived from the YTHDF2 protein were consistently the most prominent feature in SLE patients irrespective of disease activity as defined by SLEDAI score. To precisely map the YTHDF2 epitope, a phage-based alanine scanning library (PhASER) was constructed, in which each individual residue of the YTHDF2-enriched peptide was systematically substituted with alanine. Complementary sliding-window libraries were constructed by tiling six consecutive alanine substitutions across the full 49aa peptide in 3-residue increments, and a parallel series of sliding early stop codon libraries was used to define the contribution of C-terminal sequence to antibody binding. Together, these three orthogonal mutagenesis strategies identified a core set of residues required for antibody recognition, defining a shared core motif QxxPxNAY, flanked by single residues upstream and downstream of the core. This epitope was remarkably stereotypical across all 25 SLE patients with positive YTHDF2 enrichment antibodies tested (**Fig. 4A, Suppl. Fig. 4A**). Interestingly, when inspecting amino acid sequences of the eight antigens co-occurring with YTHDF2 (**Fig. 2B**), three of these (CHD9, SDK1, P4HA2) shared the same identified Px[S/N]AY motif, suggesting convergence on a common motif across this subset of SLE-associated reactivities (**Suppl. Fig. 4B**).

**Figure 4.**
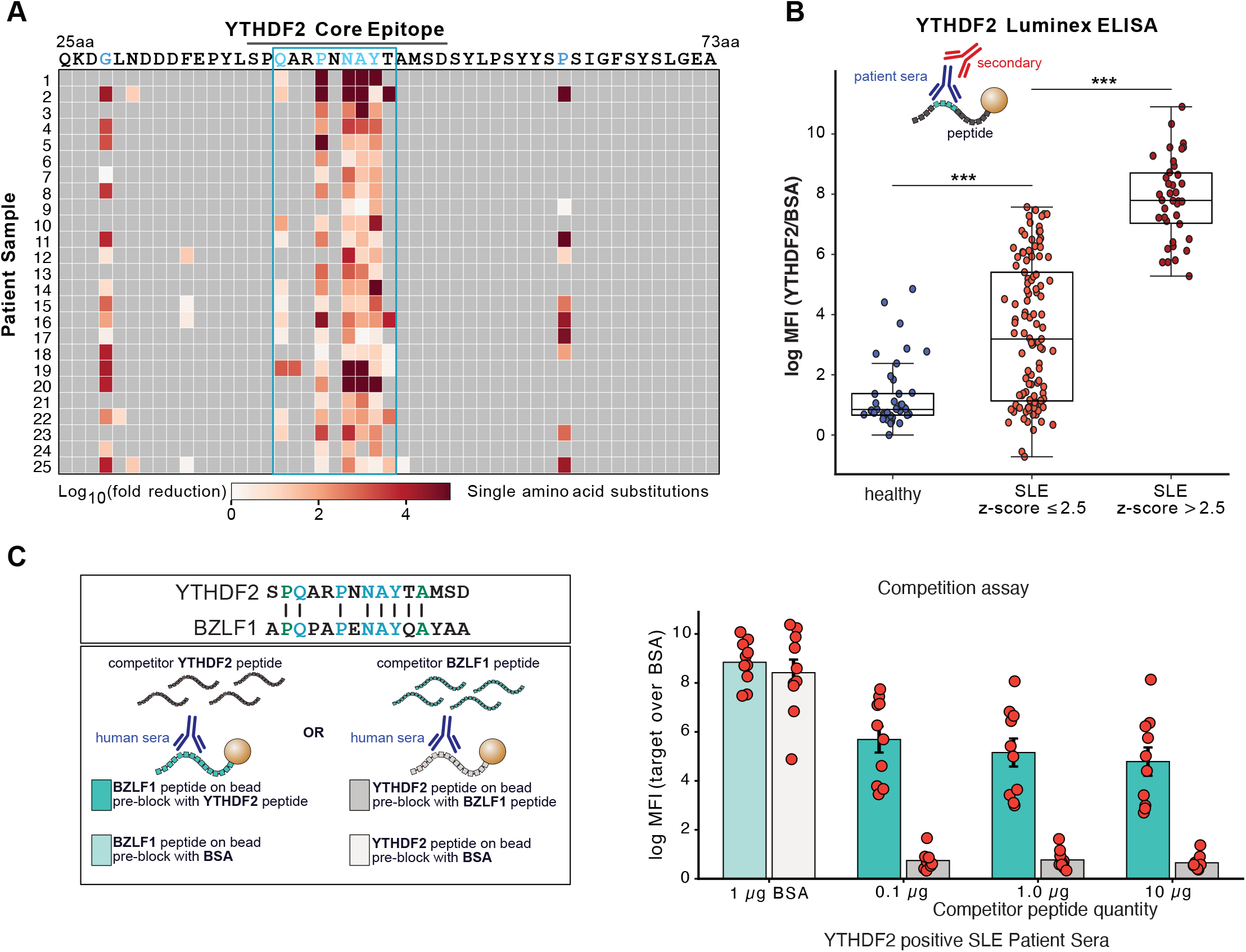
Identification of YTHDF2 epitope reveals homology with EBV BZLF1. A. Fine epitope mapping of the enriched YTHDF2 peptide by single-residue alanine scanning. Heatmaps depict SLE plasma reactivity to peptides containing individual alanine substitutions at each position. Mutations within the QxxPxNAY motif abolished antibody binding, identifying this sequence as the core autoantibody epitope. B. YTHDF2 peptide reactivity measured by Luminex in healthy controls and patients with SLE stratified by YTHDF2 PhIP-seq positivity (Z-score threshold = 2.5). Each point represents an individual subject; boxplots show median and interquartile range. C. Competitive inhibition of anti-YTHDF2 antibody binding by YTHDF2 or BZLF1 peptide. Diagram of the experimental assay (left) with top box showing motif similarity between YTHDF2 and BZLF1 peptides and bottom box showing beads used with competitor peptides. Plasma from YTHDF2 autoantibody-positive patients with SLE was pre-incubated with BSA or rising concentrations of YTHDF2 or BZLF1 peptide prior to assessment of antibody binding by Luminex (right).

To orthogonally validate antibody recognition of the YTHDF2 epitope obtained through PhASER, we performed a Luminex-bead based assay measuring binding to the original YTHDF2 epitope containing 15aa peptide. Plasma from patients with SLE, stratified by PhIP-seq YTHDF2 z-score, and from healthy controls, demonstrated a stepwise increase in reactivity (**Fig. 4B**). Individuals with high PhIP-seq enrichment (z-score > 2.5) exhibited markedly elevated binding compared to both low-z-score SLE patients and healthy controls, while low-z-score SLE patients showed bimodal reactivity: some with intermediate reactivity, and some with low or baseline levels reactivity. Luminex signal intensity closely tracked with PhIP-seq enrichment, supporting concordance between these independent assays and confirming that the identified epitope corresponds to a *bona fide* antibody target in vivo. Notably, measurable Luminex reactivity was observed in around half of SLE samples with PhIP-seq enrichment below threshold of z-score of 2.5, suggesting that the bead-based assay may capture lower-abundance antibody responses not fully detected by phage display (**Suppl. Fig 4C**).

Autoreactivity patterns in this assay may reflect loss of tolerance to self-antigens or they may reflect cross-reactivity with foreign antigens. The high-resolution epitope mapping of the anti-YTHDF2 autoreactivity enabled us to search amongst common human infectious agents for possible matches. Using PROSITE, we identified a region of high similarity between the YTHDF2 epitope and the EBV immediate-early lytic switch protein BZLF1 / ZEBRA (**Fig. 4C**), raising the possibility of molecular mimicry or cross-reactivity. To test whether this reactivity reflected cross-recognition of EBV antigens, we performed competitive binding assays using Luminex beads. Plasma from SLE patients, stratified by YTHDF2 autoantibody status, and from healthy controls were pre-incubated with BZLF1 or YTHDF2 peptides (or control BSA), followed by assessment of cross-reactive binding to Luminex beads (YTHDF2 blocking with BZLF1 readout; BZLF1 blocking with YTHDF2 readout). Plasma from YTHDF2 autoantibody-positive patients exhibited robust binding to BZLF1-coated beads, whereas healthy controls showed minimal reactivity (**Fig. 4C, Suppl. Fig. 4D**). Pre-incubation with YTHDF2 peptide reduced binding to BZLF1, consistent with recognition of a shared epitope. Conversely, pre-blocking with BZLF1 peptide completely abrogated binding to YTHDF2, even at low peptide concentrations (**Fig. 4C**, 0.1 μg), indicating high-affinity cross-reactivity. Together, these findings suggest the autoreactive enrichment of the YTHDF2 we observed in SLE patients is driven by antibody recognition directed against lytic EBV, given the critical role of BZLF1 during the lytic stage of the EBV lifecycle. This cross-reactivity was not explained by differences in conventional EBV serostatus, as anti-EBNA1 IgG and viral capsid antigen (VCA) IgG levels were comparable across YTHDF2 autoantibody positive and negative SLE patients as well as healthy controls (**Suppl. Fig. 4E**). Thus, the observed reactivity likely reflects a selective, epitope-level response against BZLF1 rather than broader anti-EBV immunity differences. Notably, YTHDF2 autoantibody positivity did not differ by immunosuppression status among SLE patients (17.8% vs. 19.3%; OR 0.91, p = 0.72), arguing against immunosuppression-associated EBV reactivation as the primary driver of this reactivity.

### A compact peptide panel recapitulates the SLE-associated autoreactive signature

To determine whether the complex SLE-associated autoreactive signature identified by PhIP-seq could be captured using a simplified and scalable assay format, we selected 10 of the highest-performing peptides for representation on Luminex beads (**Fig 5A**). An elastic-net logistic regression model was trained using the Luminex measurements of peptide reactivity from a training cohort comprising 149 SLE patient samples and 40 healthy controls (cohort #1), generating a weighted composite score termed the Lupus Autoantibody Index (LAI). In the training cohort, the model achieved a cross-validated AUC of 0.95. Analysis of out-of-fold probabilities identified a prespecified optimal LAI classification threshold of 0.773 (minimizing distance to (0,1)), yielding 87.3% sensitivity and 92.5% specificity in training. This prespecified model and threshold were subsequently applied, without further tuning or parameter fitting, to 192 samples from independent validation cohort #2 comprising 142 SLE patient samples and 50 healthy controls. In this validation cohort, the frozen LAI demonstrated high discriminatory performance with an AUC of 0.95 (95% CI: 0.92–0.98). Applying the prespecified 0.773 cut-off yielded an 88.0% sensitivity (125/142; 95% CI: 82.1–93.2%) and a 96.0% specificity (48/50; 95% CI: 86.3–99.5%), confirming robust classifier transfer across independent cohorts relative to healthy individuals. Notably, the index integrated information across multiple autoreactivities, including BZLF1, DIDO1, LOC100996413 (also known as XM_047446522), DDX17, and SNRNP70, capturing a broader humoral immune signature than any individual antigen alone. Together, these findings demonstrate that a complex PhIP-seq-derived autoantibody signature can be compressed into a compact and scalable assay format.

**Figure 5.**
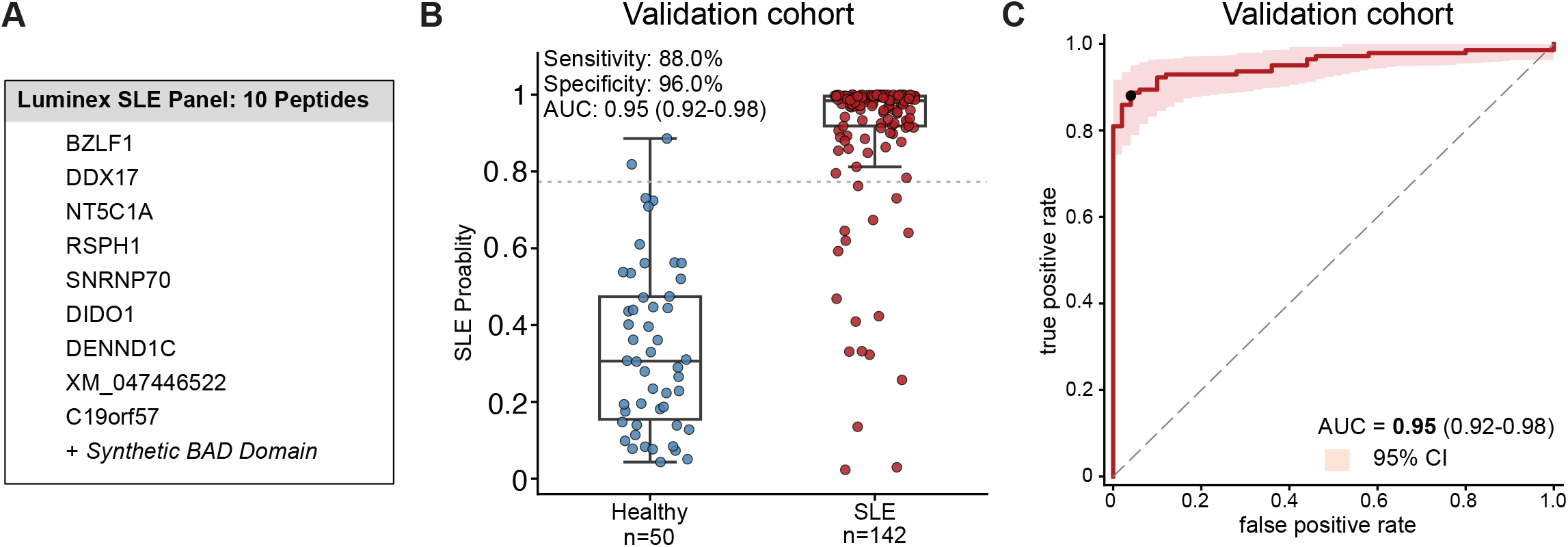
Performance of a 10-peptide Luminex panel for classification of SLE. A. Ten-peptide human autoantigens panel prioritized from PhIP-seq screening, as well as synthetic BAD domain and YTHDF2 cross-reactive peptide BZLF1. B. Distribution of the Lupus Autoantibody Index (LAI) in an independent validation cohort. The Lupus Autoantibody Index derived from an elastic-net model integrating reactivity to the ten-peptide SLE panel measured by Luminex in 189 samples from training cohort #1 was applied without refitting to the Luminex 10 peptide-reactivity measurements from 192 samples from validation cohort #2. Each point represents a sample; boxplots show median and interquartile range. The dashed line indicates the prespecified classification threshold of 0.773, derived from training cohort #1. C. Receiver operating characteristic (ROC) analysis of the LAI score in 189 samples from validation cohort #2. Black point indicates the prespecified operating point derived from training cohort #1. Shading is a pointwise 95% band from bootstrap resampling of donors.

## Discussion

A central feature of SLE is the emergence of self-reactive adaptive immune responses years before the onset of the disease, with progressive diversification of humoral immunity against self-antigens^43,44^. Although multiple autoantibody specificities have been described in lupus, the mechanisms that initiate and sustain this remarkably broad autoreactive repertoire remain incompletely understood. In this study, unbiased proteome-scale serological profiling using tiled 49aa peptides revealed extensive enrichment of shared antibody responses in patients with SLE compared with controls, identifying a previously unappreciated spectrum of autoreactivity against both viral and host antigens. These data complement existing SLE associated autoantigens that were discovered using whole proteins. Rather than appearing as unrelated autoreactivities, the SLE-associated autoreactivities discovered here clustered around proteins involved in RNA metabolism including within the spliceosome, transcriptional regulation, nuclear pore related, nucleic acid binding, and cell death. The convergence of these antigenic targets suggests that lupus autoimmunity represents the immune recognition of interconnected molecular systems rather than individual proteins. Our orthogonal validation, using a compact panel of only 10 peptide targets, robustly distinguished SLE patients from healthy individuals, underscoring the degree of convergence within this autoreactive signature. A limitation of this approach is that the compact panel was only used to classify relative to healthy samples, and it will be of interest to determine whether related interferon-driven or autoimmune conditions share features of this signature.

Beyond related molecular systems, we find that the autoreactivity of the most SLE-associated peptides is driven by highly similar features, including an enrichment of BAD proteins and sequences predicted to lie within intrinsically disordered regions. Although these proteins often exhibit limited sequence homology, they share the capacity to interact with nucleic acids and to assemble into large macromolecular complexes involved in RNA metabolism. The association of these proteins with nucleic acids may further contribute to their immunogenicity by coupling B-cell recognition of RNA-binding proteins to internalization of associated RNA and activation of endosomal TLR7 and type I interferon pathways.

This concept may help explain a longstanding feature of lupus serology: the tendency of autoantibody responses to spread across groups of RNA-binding proteins despite limited primary sequence similarity. Rather than being driven exclusively by shared peptide sequence, autoreactive B cell selection may be influenced by common biophysical properties and participation in higher-order ribonucleoprotein structures.

Although direct evidence linking phase-separated assemblies to immune-complex architecture in lupus remains limited, it is possible that these properties may amplify pathogenicity by promoting the formation of large immune complexes. Antibody engagement of multivalent, condensate-associated proteins could facilitate cross-linking and aggregation, enhancing immune complex deposition and downstream inflammatory signaling. Similar mechanisms have been proposed in neurodegenerative diseases, where aggregation-prone domains contribute to pathological protein assemblies^45^.

Among the RBPs, peptides derived from the m6a-reader protein YTHDF2 were among the most significant differentiators between SLE and healthy controls. High resolution mapping of the anti-YTHDF2 epitope, followed by a search of human viral sequences surfaced a close match to BZLF1 (aka Zta/ZEBRA/EB1), one of the key lytic switch immediate-early transcription factors crucial for EBV reactivation through binding of Zta-responsive elements. Competition assays demonstrated reciprocal cross-reactivity between these antigens, providing direct evidence of epitope-level molecular mimicry. Importantly, this reactivity is not explained by differences in conventional EBV serostatus, indicating that it reflects selective recognition of specific viral epitopes rather than generalized antiviral immunity. BZLF1 is a well-studied bZIP transcription factor with a long intrinsically disordered region, and its presence as a soluble factor in serum, perhaps due to passive release after cell lysis or through exosome secretion, has been considered as a biomarker for complications related to EBV, such as post-transplant lymphoproliferative disease^46^. The relationship between EBV and lupus has been suspected for decades, supported by epidemiological studies demonstrating increased EBV seropositivity in pediatric patients as compared to controls^47^, elevated latent and lytic gene expression^48^, and cross-reactivity between EBNA1 and Sm and Ro antigens^49^. More recently, host genetic variations were associated with persistent EBV DNAemia in multiple autoimmune diseases including SLE, at a population-scale level^50^. EBV+ B cell subsets in SLE were enriched in the CD27^+^CD21^low^ memory B cell, an activated memory pathologically expanded in SLE^19^.

The identification of cross-reactivity between YTHDF2 and the EBV immediate-early protein BZLF1 raises the possibility that this autoreactive response may arise in the context of EBV lytic reactivation. Differentiation of memory B cells into plasmablasts can induce expression of the transcription factors XBP1 and BLIMP1, which can directly activate BZLF1 and trigger EBV lytic reactivation^51^. In this context, transient expression of BZLF1 may provide a source of antigenic stimulation that preferentially selects for B cells recognizing structurally similar host epitopes, including YTHDF2.

EBV lytic reactivation is initiated by the immediate-early transcription factor BZLF1 and is accompanied by extensive remodeling of host transcriptional and chromatin programs, activation of DNA damage response pathways including ATM and ATR signaling, metabolic and oxidative stress responses, and engagement of apoptotic regulatory networks required for productive viral replication^52,53^. In the context of SLE, where clearance of apoptotic and secondary necrotic material is impaired, repeated cycles of EBV reactivation may generate a sustained source of extracellular viral and host nucleoprotein antigens capable of driving epitope spreading and innate immune activation, especially concerning chromatin and RNA binding proteins, and those prone to forming condensates.

Together, these results greatly expand the known range of SLE-associated, and SLE disease activity associated autoreactive peptides and their respective proteins. The landscape of SLE autoreactivity described here is complementary and largely distinct from previously characterized autoantigens based on whole proteins. However, both sets share common cellular processes, especially RNA processing. The features described here may contribute not only to our understanding of immune dysfunction in SLE, but also may provide a foundation for determining whether these autoreactivities have future utility in clinical stratification or longitudinal assessment, including with emerging B-cell depletion strategies, such as CD19 and BCMA CAR-T. Finally, this data suggests a model in which SLE-associated humoral autoreactivity emerges or is driven by the intersection of EBV viral mimicry, epitope convergence, and the intrinsic biophysical properties of target antigens.

## Acknowledgments

We thank The Immune Tolerance Network, L. Cooney and L. Higdon for sharing samples from the CALIBRATE and ACCESS trials. We thank the patients for their participation in this study.

This work was supported by the Lupus Foundation Award, Inc (CH), the NIH grant T32AR079068 (CH), Chan Zuckerberg Biohub San Francisco (JLD, MSA, CJY), Westridge Foundation (MRW), NIH/NIMH R01MH122471 (MRW, JLD), NIAMS P30AR070155 (JY, MDA), CDC U01DP006701 (MDA, JY), CDC U01DP006700 (PI, JB), NIH/NIAID UM1AI109565 (MSA), NIAID R01AI171184 (CJY), NIAMS P30AR070155 (CJY), the Department of Defense (CJY), the Arc Research Institute (CJY), Parker Institute for Cancer Immunotherapy (CJY).

Sequencing was performed at the UCSF Center for Advanced Technology, supported by UCSF PBBR, RRP IMIA, and NIH 1S10OD028511-01 grants.

## METHODS

### Human cohorts and samples

The primary discovery cohort (cohort #1) consisted of participants enrolled in the California Lupus Epidemiology Study (CLUES), a prospective longitudinal cohort of individuals with systemic lupus erythematosus (SLE) followed at the University of California, San Francisco (UCSF). Recruitment started in 2015. The cohort is broadly representative of the demographic characteristics of SLE, including a predominance of female participants (89%; Table 1), and spans a wide spectrum of disease activity, ranging from inactive disease (Systemic Lupus Erythematosus Disease Activity Index [SLEDAI] = 0) to high disease activity across multiple systems (maximum SLEDAI = 44). SLE diagnoses were confirmed by study physicians based on fulfillment of the 1997 American College of Rheumatology (ACR) classification criteria, fulfillment of three ACR criteria together with a rheumatologist-confirmed diagnosis of SLE, or biopsy-confirmed lupus nephritis. Plasma samples and detailed clinical metadata were collected at routine longitudinal visits. An additional flare cohort enrolled participants at the time of clinically confirmed disease flares. Some individuals contributed samples to both the longitudinal and flare cohorts. In total, the discovery cohort comprised 890 plasma samples from 475 individuals with SLE. All samples were processed in parallel and randomized across experimental plates to minimize batch effects. Healthy controls included participants enrolled through the healthy control arm of the CLUES study together with healthy donors obtained from the New York Blood Center.

External validation was performed using two independent SLE cohorts. The first validation cohort (cohort #2) comprised participants enrolled in the Immune Tolerance Network (ITN) ACCESS^54^ and CALIBRATE^55^ clinical trials (n = 123 and n=38 SLE patients respectively), followed longitudinally, with 297 total SLE samples, along with 91 healthy controls from the HBDB biobank. The second validation cohort (cohort #3) included an independent set of samples from SLE patients enrolled at the New York University (NYU) comprising 81 plasma samples from 59 individuals with SLE. 48 healthy controls were included in cohort #3, of who 46 had also been processed as part of cohort #1. To prevent information leakage between cohorts, these overlapping HCs were excluded from cohort #1 for all analyses involving prediction of cohort #3.

All studies were approved by the Institutional Review Boards of the participating institutions, and all participants provided written informed consent prior to enrollment.

### PhIP-seq with human peptidome

We performed PhIP-seq as previously described following established protocols (protocol.io^56^). Briefly, we used a human peptidome T7 phage display library comprising 731,724 unique bacteriophage, each expressing a 49–amino acid peptide with 25–amino acid tiling overlap across the human proteome.

Deep-well plates were pre-blocked overnight at 4°C with 3% bovine serum albumin (BSA) in TBST to minimize nonspecific binding. For each reaction, 1 μl of human plasma diluted 1:1 in storage buffer was incubated with 500 μl of freshly amplified phage library overnight at 4°C with gentle rocking. Antibody-phage complexes were captured using protein A/G magnetic beads (Thermo Fisher), followed by several rounds of washing to remove unbound phage.

Bound phage were then eluted directly into log-phase Escherichia coli (BLT5403; OD□ □ □ = 0.5–0.7) and amplified by bacterial infection at 37 °C. The amplified phage pool was subsequently re-incubated with plasma from the same individual, and this cycle of immunoprecipitation and amplification was repeated for a total of three rounds to enrich antigen-specific phage. Following the final round, phage DNA was extracted, barcoded, PCR-amplified with Illumina adapters, and subjected to next-generation sequencing on an Illumina platform to a depth of approximately 1 million reads per sample.

Sequencing reads were aligned to the reference phage library. All analyses were performed at the peptide level. Peptides with fewer than 10 total reads across all samples were excluded, resulting in a final dataset of 360,920 peptides. To permit inclusion of peptides with zero observed reads across mock IP and healthy controls in downstream analyses, a pseudocount of one read was added to each peptide in each sample. Peptide enrichment was quantified as fold enrichment relative to mock immunoprecipitation controls by normalizing peptide-specific read counts to the mean read count for the same peptide across AG bead–only control samples. To account for technical variation between assay plates, batch effects were corrected using linear regression. Batch-corrected enrichment values were subsequently compared to the distribution observed in healthy controls, and peptide-specific z-scores were calculated for each peptide in each individual. A threshold of z-score ≥ 2.5 was used to define antibody reactivity. A commercial anti-GFAP polyclonal antibody was included as a positive control, and showed strong enrichment of GFAP-derived peptides, accounting for >50% of reads on average, confirming assay performance.

### Differential autoantibody enrichment analysis

To identify which individual protein targets show SLE-specific autoantibody enrichment, each peptide was tested independently for association with disease status by univariate logistic regression. The model for each peptide was logit(P(SLE)) = β0 + β1 × donor-mean z-score, where z-scores were computed from the reactivity z-score layer (z relative to the healthy-control mean and standard deviation) and averaged within each donor. Each peptide was modeled independently, with no regularization and no joint optimization across peptides. The Wald z-statistic (β1/SE(β1)) served as the per-peptide test statistic. Models were fit by iteratively reweighted least squares.

Protein-level significance was derived by applying Fisher’s combined test to the two-sided p-values of all peptides mapping to each protein, giving a combined statistic equal to −2 × Σ log(p□). The two-sided p-value for each peptide was obtained from its Wald z-statistic. An empirical null distribution was constructed by permuting the SLE and healthy-control donor labels 100,000 times and repeating the full per-peptide regression and protein-level aggregation under each permuted labeling. Because permutation preserves the correlation among peptides within a protein while breaking the association with disease status, the protein-level test does not assume independence among peptides. For each protein, the empirical p-value was (b + 1) / (M + 1), where b is the number of permutations whose Fisher statistic equaled or exceeded the observed value and M is the total number of permutations. Multiple testing correction across proteins used the Benjamini-Hochberg procedure at a false discovery rate of 0.1.

The volcano plot shows on the x-axis the signed Wald z-statistic of the most strongly associated peptide per protein, defined as the peptide with the smallest two-sided p-value, taken from the observed logistic regression. Positive values indicate higher reactivity in SLE and negative values indicate higher reactivity in healthy controls. The y-axis shows −log10 of the protein-level permutation p-value. The direction of enrichment for each significant protein was assigned from the sign of this peptide. For significant SLE-enriched proteins, point color encodes the fraction of SLE donors with a reactivity z-score of at least 4.5 at that peptide, capped at 10 percent. Significant healthy-control-enriched proteins are shown in a single color and non-significant proteins in gray.

The same pipeline was applied to patients with high disease activity (SLEDAI ≥ 11) versus patients with no disease activity (SLEDAI = 0).

### Autoantibody diagnostic classifier

To test whether serum antibody reactivity discriminates SLE from healthy control, an elastic-net logistic regression classifier was trained on all peptides jointly, estimating every peptide coefficient simultaneously under combined L1 and L2 regularization. Class weights were set in inverse proportion to class frequency within each training set. The classifier used C = 1.0, an L1 fraction of 0.5, the SAGA solver, a tolerance of 0.001, and 100 iterations.

Internal performance was estimated using the full UCSF cohort of (890 SLE and 208 healthy controls). Five-fold cross-validation was performed with folds grouped by donor, ensuring that all longitudinal samples from an individual were assigned to the same fold and that no donor appeared in both training and testing sets. Cross-validation was repeated across 20 random seeds, resulting in 100 model fits. Area under the receiver operating characteristic curve (AUROC) was computed for each held-out fold and summarized as the mean across the 100 fits with the 2.5^th^ to 97.5^th^ percentile range.

External validation was performed using the two aforementioned independent cohorts. Of note, while all the samples in validation cohort #2 were independent from cohort #1, for validation cohort #3 (NYU), a subset of healthy control samples from cohort #1 was reprocessed in parallel with the NYU samples and used as the healthy comparison group. To prevent overlap between training and validation datasets, these healthy controls were excluded from model training. The classifier was then retrained on the reduced UCSF dataset (890 SLE and 162 healthy controls), and the resulting 100 models were applied to the external validation cohorts.

### Feature-count ablation

To characterize how performance scales with the number of input peptides, peptides were ranked by the magnitude of their median coefficient across the 100 full-feature models, among peptides with a nonzero coefficient in at least 90% of the fits. For each of 19 feature-counts, every integer from 1 to 10 and then 20, 30, 40, 50, 100, 200, 300, 400, and 500, a new set of elastic-net models was trained on the reduced UCSF dataset restricted to those peptides, using the settings above with up to 200 iterations, and applied to Cohort #2 and Cohort #3. Area under the curve was summarized across the 100 model fits for each feature count and cohort.

### Patient-level antibody enrichment heatmaps

Sample-level reactivity was displayed as a blockwise heatmap. The rows are the most significantly enriched peptide for each of the top SLE-enriched proteins from the univariate permutation analysis, taking proteins with gene-level FDR below 0.1 and ranking them by signed Wald z-statistic, with one peptide per protein chosen as the peptide with the smallest two-sided p-value. The 20 proteins shown were taken from the top of this ranking. Columns are individual serum samples from the cohorts #1, #2 and #3, split into SLE and healthy controls and grouped by cohorts within each. Each cell shows the reactivity z-score for one sample and one peptide, computed relative to the healthy-control mean and standard deviation. The colormap shows z-scores at or below 2.5 in white and scales through a red gradient to a maximum at 10. A track above the heatmap shows the number of the displayed peptides reaching a z-score of at least 2.5 in each sample.

### Peptide co-enrichment analysis

Pearson correlations of reactivity z-scores were computed across all 1,098 UCSF samples for one representative peptide from each of the 50 highest-ranked SLE-enriched proteins, taking proteins with protein-level FDR below 0.1 and ranking by signed Wald z-statistic. Within each protein, the peptide with the smallest two-sided individual p-value was used. The resulting 50 × 50 Pearson correlation matrix was displayed as a heatmap with rows and columns ordered by hierarchical clustering.

For each of the two protein clusters identified by hierarchical clustering of the 50-peptide correlation matrix, within-cluster reactivity concordance was summarized as the mean pairwise Pearson correlation of donor-level reactivity z-scores across all cluster members. Because Pearson correlation coefficients are not linearly averageable, pairwise correlations were first Fisher z-transformed (z = arctanh(r)), averaged, and back-transformed (r = tanh(z□)) to obtain the reported mean. For a cluster of size k, 100,000 random k-protein subsets were drawn without replacement from the same 50-protein candidate pool, the same Fisher z-averaged statistic was computed for each draw, and an empirical p-value was calculated as the proportion of random draws whose value equaled or exceeded the observed cluster’s, with a +1 continuity correction applied to both the denominator and numerator.

### RNA-binding protein enrichment

Human RNA-binding proteins (n = 2,481) were obtained from Trendel *et al*.^57^, of which 2,389 were present in the 19,542-protein set tested. To assess whether RNA-binding proteins concentrate at the extremes of the differential enrichment ranking, the fraction of RNA-binding proteins among the top K proteins was computed for both tails of the ranking, the SLE-enriched tail with proteins ordered by descending signed Wald z-statistic and the healthy-control-enriched tail with proteins ordered by ascending z-statistic. This fraction was evaluated (1) across the full ranking, at 120 log-spaced values of K from 50 to 19,542, and (2) restricted to the top of the ranking, at 80 evenly spaced values of K from 20 to 500. For each tail, an empirical null was generated by permuting RNA-binding protein membership across proteins 100,000 times while holding the ranking fixed, and recomputing the fraction curve on each permutation; the 2.5th and 97.5th percentile fraction at each K define the shown 95% band. The overall prevalence of RNA-binding proteins in the protein set (p□ = 0.122) is shown as a baseline.

To test for enrichment, each fraction curve, including observed and each of the 100,000 permuted nulls, was reduced to a single statistic. The area between the curve and the baseline prevalence, integrated over K by the trapezoidal rule and normalized by the K range, with values below baseline set to zero so that only excess concentration (not depletion) contributes. An empirical p-value was computed as the proportion of permuted-null statistics greater than or equal to the observed statistic, with a +1 continuity correction applied to numerator and denominator. Because RNA-binding proteins are represented by more peptides per protein on average (median 14 vs. 10 peptides/protein; Mann-Whitney p = 2.2×10□²□), evaluating both tails under the same null distinguishes enrichment specific to the SLE-enriched direction from a generic effect of peptide count per protein.

To test whether the RNA-binding protein enrichment above was disproportionately driven by XIST-associated RNA-binding proteins, XIST-interacting proteins (n = 129) were intersected with the RNA-binding protein set (103 genes annotated as both). Restricted to RNA-binding proteins, a logistic regression predicted SLE-significance (BH-FDR q ≤ 0.1) from XIST-associated membership, controlling for log(1 + peptides per gene) to account for the peptide-count effect noted above. XIST-associated membership was not a significant predictor of SLE-enrichment significance among RNA-binding proteins (odds ratio = 1.44, 95% CI 0.70– 2.98, p = 0.32).

### Disease activity classifier

A second elastic-net classifier was trained with the same peptide input, hyperparameters, class weighting, and donor-grouped cross-validation design as the SLE versus healthy-control classifier. The disease activity classifier separates UCSF donors with high disease activity (SLEDAI at least 11) from donors with no disease activity (SLEDAI 0). Threshold of 11 was chosen according to prior studies^58,59^. Samples with intermediate SLEDAI were not used, and donors contributing samples to both groups were removed. The 100 trained models were applied without retraining to the two independent validation cohorts, restricted to lupus samples with SLEDAI at least 11 or 0.

### pI and charge calculations

To test charges of peptides, we computed the isoelectric point (pI) in two ways: for the full 49-residue peptide, and as the maximum pI across all overlapping 12-residue windows tiled along the sequence. Two companion statistics were computed alongside pI using the same framework: the fraction of lysine-plus-arginine residues (K+R fraction) and the net charge at physiological pH (7.4). For each statistic, we built a null distribution from 100,000 random 7-residue peptides drawn from the full peptidome, computing the identical statistic for each draw.

### Condensate and BAD domain enrichment

Condensate-forming propensity scores were obtained from the PICNIC database. Among proteins reaching FDR below 0.1 with positive z-statistic in the high versus no disease-activity analysis (N=111), the median PICNIC score was compared to an empirical null of 100,000 same-size random draws (without replacement) from the full PICNIC-scored gene library. An empirical p-value was computed as the proportion of null draws with a median PICNIC score at or above the observed value.

### PhASER

PhASER library containing deep mutational scanning of some of the SLE-enriched antigens were created with 1. Mutations of each amino acid in the peptide sequence to an alanine (single alanine scan), and if peptide were already an alanine, mutation of the alanine to a glycine; 2. Mutations of each amino acid in the peptide sequence to a stop codon; 3. Mutations of 6 amino acid segments to alanine with sliding window of 3 aminoacids overlapping. Libraries were immunoprecipitated with patients, controls and mock plasma as described above.

PhIP-seq read counts for each peptide variant were normalized against the input library by converting counts to within-sample frequencies, dividing by corresponding input library frequencies, and log□-transforming the resulting relative abundances. A pseudocount of 0.5 was added prior to normalization. Samples were considered reactive against a given wild-type peptide if their mean log□ enrichment across all variants exceeded 1, corresponding to at least two-fold enrichment over input.

### SLBA

We performed SLBA as reported previously^60,61^. Briefly, the nucleic acid for our top enriched SNRNP70 peptide (sequence 5’-KGRDRDRERRRSHRSERERRRDRDRDRDRDREHKRGERGSERGRDEARG-3’) was inserted into a split luciferase construct containing a terminal HiBiT tag and synthesized as DNA oligomers (IDT). Progressive mutations of the basic acidic residues were made in addition to the original sequence and synthesized as DNA oligomers, with 10%, 20%, 40%, 60%, 80%, 90% and 100% change of basic and acidic residues towards alanine. Constructs were amplified by PCR using 5′-AAGCAGAGCTCGTTTAGTGAACCGTCAGA-3′ and 5′-GGCCGGCCGTTTAAACGCTGATCTT-3′ primer pair. Unpurified PCR product was used as input to rabbit reticulocyte transcription translation system (Promega), and Nano-Glo HiBiT Lytic Detection System (Promega cat no. N3040) was used to measure relative luciferase units (RLU) of translated peptides in a luminometer. Peptides were normalized to 5e6 RLU input, incubated overnight with patient sera, and immunoprecipitated with protein A and protein G Sepharose beads (MilliporeSigma). After thoroughly washing beads with SLBA buffer (0.15 M NaCl, 0.02 M Tris-HCl, pH 7.4, 1% wt/vol sodium azide, 1% wt/vol bovine serum albumin, and 0.15% vol/vol Tween-20), luminescence remaining on beads was measured using Nano-Glo HiBiT Lytic Detection System (Promega cat no. N3040) in a luminometer. Anti-HiBiT antibody (Promega) was used as a positive control for each peptide. Fold change of RLU over peptide only background was calculated for each patient.

### Identification of proteins sharing the QxxPxNAY motif

To identify proteins containing a sequence motif shared with YTHDF2, the QxxPxNAY motif was queried using ScanProsite (ExPASy; https://prosite.expasy.org/scanprosite/) with the “Submit MOTIFS to scan them against a PROTEIN sequence database” option. The motif was entered as a user-defined pattern and searched against the UniProtKB protein sequence database. ScanProsite permits user-defined amino-acid patterns in which “x” represents any amino acid. The search identified 22 protein matches. The top hit corresponded to a protein from *Pyrobaculum aerophilum*, an archaeal species not known to infect humans. Notably, hits 2 to 4 corresponded to the BZLF1 protein of human herpesvirus 4 (HHV-4/EBV). Four additional hits corresponded to YTHDF2 orthologs containing the shared motif across four mammalian species, including *Homo sapiens*, *Mus musculus*, *Macaca fascicularis*, and *Bos taurus*.

### Luminex assay

Peptides containing the relevant motif from YTHDF2, BZLF1, RSPH1, DDX17 found via PhASER, or full 49 aminoacids peptides recognized by PhIP-seq from NT5C1A, FIP1L1, SNRNP70, C19orf57, NPIPB11, DIDO1, LOC100996413, DENND1C, as well as consensus motif derived from BAD domains (‘BAD’) were synthesized (Genscript, LifeTein) and conjugated to BSA via SMCC coupling to cysteines on BSA at a 1:1 final mass ratio. Spectrally distinct Luminex beads were each conjugated to an individual BSA-peptide. A control bead population was made by conjugating BSA only. Each bead conjugation was performed at a concentration of 5□μg of protein per 1 million beads in a 0.5-ml reaction volume. Conjugation was done via an EDC/sulfo–N-hydroxysuccinimide coupling strategy to free amines using an Antibody Coupling Kit following the manufacturer’s instructions (Luminex, 40-50016).

All serological analyses were performed in duplicate, and beads were pooled on the day of use. Thawed plasma samples were diluted 1:250 in PBS + 0.05% Tween 20 (PBST) containing 2% non-fat milk and mixed with pooled peptide-coated beads (2,000–2,500 beads per peptide). Samples were incubated for 1□h at room temperature at 250 r.p.m, washed three times with PBST and then stained with 1:2,000 phycoerythrin-conjugated anti-human IgG Fc antibody (BioLegend, 637310) in PBST for 30□min at room temperature. Beads were washed twice with PBST and analyzed in a 96-well plate format on a Luminex LX 200 cytometer. For each sample, peptide MFI values were normalized to the BSA-only control (peptide MFI ÷ BSA MFI) and averaged across duplicate wells.

### Luminex autoantibody index

To derive a composite serological score from the Luminex assay, we selected ten peptide reactivities corresponding to the PhIP-seq–prioritized lupus-associated epitopes that showed significant enrichment in SLE and/or in high disease activity. An elastic-net logistic regression model was trained on these ten Luminex-derived features to distinguish patients with SLE from healthy controls, using a discovery cohort of 149 SLE patients and 40 healthy controls (cohort #1). Model tuning was performed via 5-fold cross-validation within the training cohort to select the elastic-net mixing proportion and regularization parameter. The Lupus Autoantibody Index (LAI) was defined as the model’s predicted probability of SLE, obtained by applying the logistic function to the linear predictor (the sum of standardized peptide reactivities weighted by the model coefficients, plus the intercept). The LAI therefore ranges from 0 to 1, with higher values indicating a more lupus-like autoantibody profile. An operating threshold of 0.773 was prespecified on out-of-fold training probabilities by minimizing the Euclidean distance to the ideal point (0,1) on the ROC curve. The fully fitted pipeline (feature scalers, model weights, and operating threshold) was frozen and applied without refitting or local recalibration to an independent validation cohort of 142 SLE samples and 50 healthy controls (Cohort #2, n = 192; 163 unique donors). Primary classification metrics were evaluated against the prespecified threshold, reporting exact binomial 95% confidence intervals for specificity and donor-level bootstrapped 95% confidence intervals for sensitivity and AUC to account for repeated donor sampling.

### BZLF1 / YTHDF2 competition assay

To assess cross-reactivity between BZLF1 and YTHDF2 peptides, competition assays were performed using the Luminex platform described above. BZLF1 peptide of the same size as the YTHDF2 peptide and containing the similar motif was synthesized (Lifetein). Plasma samples diluted 1:250 in PBST-2% non-fat milk were pre-incubated for 1 hour at room temperature with either soluble BSA-conjugated BZLF1 or BSA-conjugated YTHDF2 peptide at final amounts of 0.1, 1, or 10□μg for each sample, or with BSA alone. Following pre-incubation, plasma samples blocked with BZLF1 peptide were added to YTHDF2-conjugated beads, whereas plasma samples blocked with YTHDF2 peptide were added to BZLF1-conjugated beads and gently rocked at 250rpm for an hour followed by 3 washes and staining with PE anti-human IgG Fc antibody and analyzed as above.

---

**Supplemental Figure 1.**
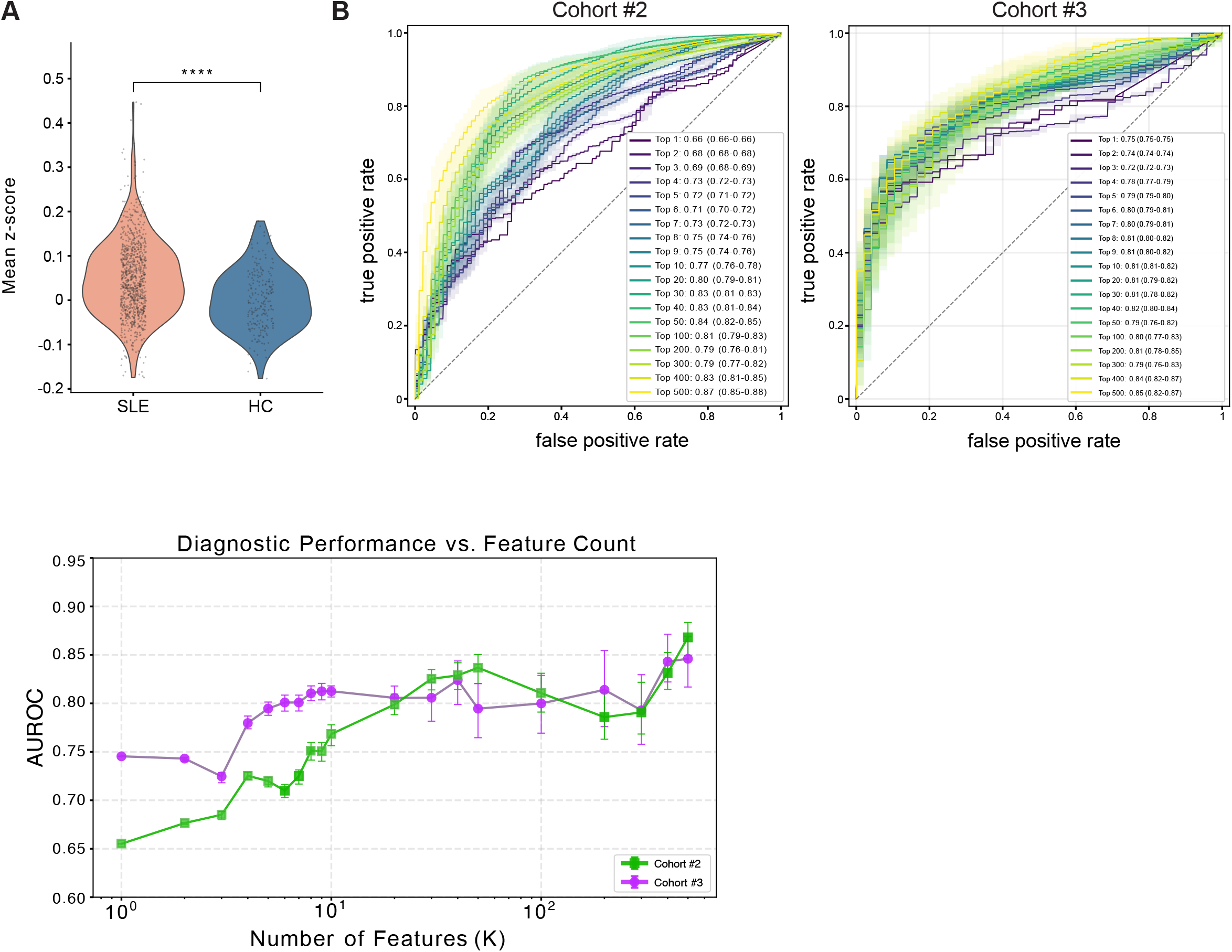
A. Overall magnitude of autoreactivity in SLE as compared to HC across all 360,920 peptides with more than 10 reads across the whole study population (****, p<0.0001). B. Receiver operating characteristic (ROC) curves showing the diagnostic performance of the top predictive peptides identified in the training cohort and evaluated in the independent validation cohorts #2 and #3. Curves represent models incorporating increasing numbers of top-ranked predictive peptides, as indicated. C. Diagnostic performance (AUROC) in validation cohorts #2 and #3 as a function of the number of top-ranked predictive peptides included in the model. Error bars indicate 95% confidence intervals.

**Supplemental Figure 3.**
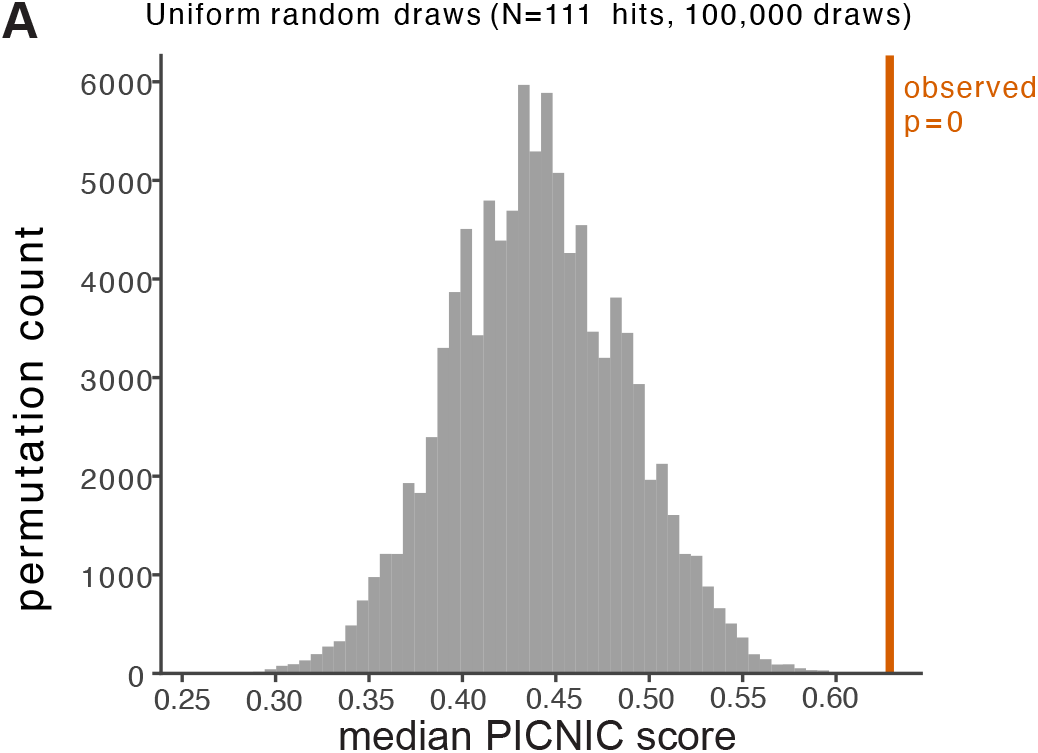
Proteins enriched in high disease activity exhibit increased predicted condensate-forming propensity. A. Median PICNIC score from 100,000 random draws of 111 proteins from the background set of 17,258 proteins (null distribution – gray) compared with observed median PICNIC score (orange) corresponding to the 111 proteins significantly enriched in patients with high versus no disease activity (*p* < 1 × 10⁻ □).

**Supplemental Figure 4.**
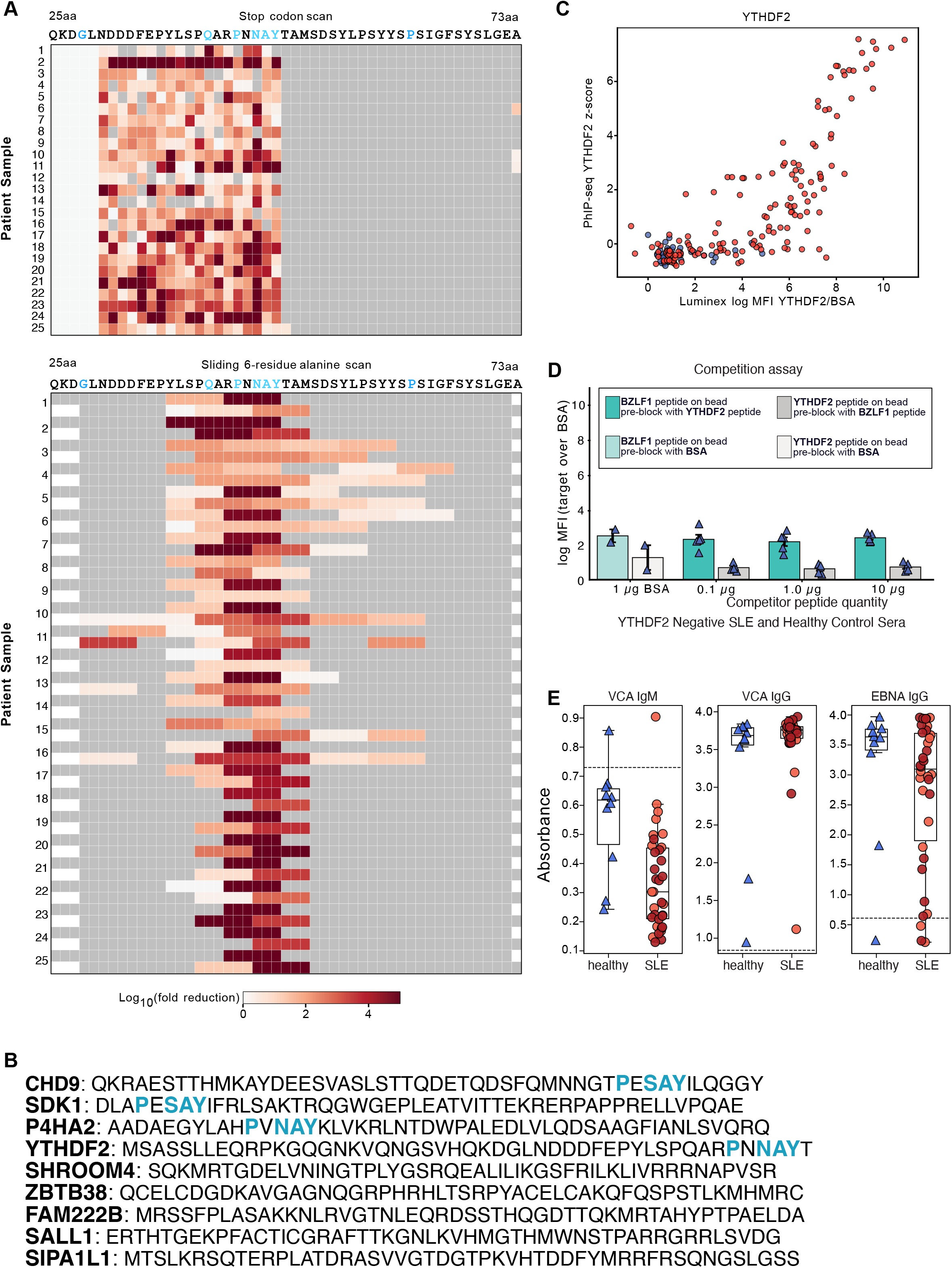
A. Epitope mapping of YTHDF2 autoantibody reactivity in 25 SLE patient samples using stop-codon scanning (top) and a sliding six-residue alanine scan (bottom). Stop codon scan shows absence of reactivity until full motif is exposed. Sliding 6 residue alanine scan shows windows around the motif is critical. Heatmaps show the log10 fold reduction in antibody reactivity relative to the unmodified peptide, identifying a shared region required for antibody binding centered on the QxxPxNAY motif. B. Sequence comparison of the 8 protein targets co-occurring with YTHDF2, showing some with related sequence motifs, with motif residues highlighted in blue. C. Correlation between YTHDF2 antibody reactivity measured by PhIP-seq (z-score) and independent YTHDF2 Luminex assay (log MFI YTHDF2/BSA). Points represent individual samples. D. Plasma from YTHDF2 autoantibody-negative patients with SLE or healthy controls was pre-incubated with BSA or rising concentrations of YTHDF2 or BZLF1 peptide prior to assessment of antibody binding by Luminex. E. EBV serologies measured by ELISA for viral capsid antigen (VCA) IgM, VCA IgG, and EBV nuclear antigen (EBNA) IgG in healthy controls and patients with SLE. Dashed horizontal lines indicate assay-specific thresholds for seropositivity.

**Supplemental Figure 5.**
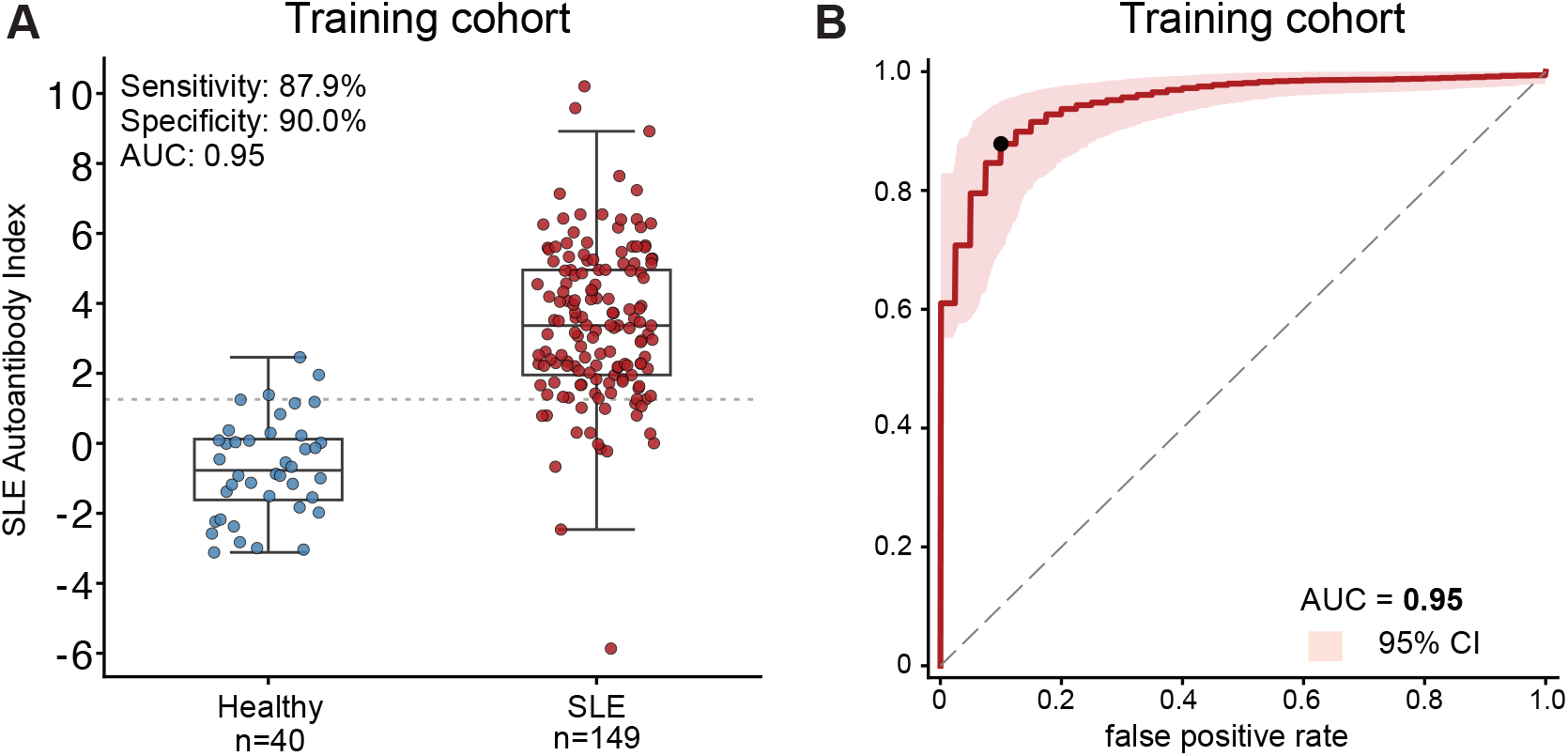
A. Distribution of the Lupus Autoantibody Index in healthy controls and SLE patients in 189 samples from training cohort #1. Scores represent the average out-of-fold probability for each sample across 20 donor-blocked cross-validation repeats. The dashed line indicates the optimal classification threshold (0.773), defined as the point of minimum distance to (0, 1) on the ROC from training cohort #1. Each point represents a sample; boxplots show median and interquartile range. B. Cross-validated ROC analysis of a 10-peptide Luminex autoantibody signature for classification of SLE versus healthy controls in 189 samples from training cohort #1, computed from the same out-of-fold probabilities as A. Black point indicates minimum distance to (0,1).

